# Can colony size of honeybees (*Apis mellifera*) be used as predictor for colony losses due to *Varroa destructor* during winter?

**DOI:** 10.1101/2020.06.22.164525

**Authors:** Coby van Dooremalen, Frank van Langevelde

## Abstract

For more than three decades, honeybee colonies (*Apis mellifera*) experience high losses during winter, and these losses are still continuing. It is crucial that beekeepers monitor their colonies closely and anticipate losses early enough to apply mitigating actions. We tested whether colony size can be used as early predictor for potential colony losses, in particular due to the parasitic mite *Varroa destructor. V. destructor* is one of the most important causes for these losses. Such early predictor for potential *V. destructor* induced losses is especially relevant as measuring *V. destructor* load in colonies is difficult and cumbersome. During three years, we monitored colonies with high and low *V. destructor* load from July until March of the next year. We found that differences in colony size were only visible after November, even though we lost almost all colonies every winter in the group with high *V. destructor* load. In the Northern hemisphere, November is considered to be too late for beekeepers to strengthen colonies in preparation for winter. We therefore argue that early-warning signs for potential colony losses due to *V. destructor* are urgently needed to allow beekeepers preventing winter losses. We discuss the role of precision apiculture to monitor the health and productivity of honeybee colonies.

## Introduction

For more than three decades, honeybee colonies (*Apis mellifera*) experience high losses during winter, especially in the Northern hemisphere, and these losses are still continuing (VanEngelsdorp and Meixner 2010, Ryabov et al. 2017, Thoms et al. 2019). It is crucial that beekeepers can anticipate these losses early enough to apply mitigating actions. Therefore methods for early warning of potential winter losses are urgently needed (Dainat et al. 2012a, Francis et al. 2013, Van Dooremalen et al. 2018). This early warning is useful when (1) winter losses can be predicted early enough for beekeepers to anticipate or intervene, and (2) the predictors can be measured relatively easy and quick. In this study, we test whether colony size can be used as an early predictor for potential losses in honeybee colonies, as larger colonies showed larger survival chance (Döke et al. 2015, Van Dooremalen et al. 2018). We aim in particular at colony size in relation to the parasitic mite *Varroa destructor*, as this mite is considered to be one of the most important causes for the current high winter losses of honeybees in the Northern hemisphere (Aizen and Harder 2009, Ellis et al. 2010, Le Conte et al. 2010, Potts et al. 2010, Rosenkranz et al. 2010).

Especially for the temperate zone in the Northern hemisphere, early detection of potential winter losses means September the latest – well before bees enter the winter cluster and during winter preparation – as this is the last window of opportunity to strengthen or merge weak colonies before they enter the winter period (Döke et al. 2015). Strengthening (i.e., adding bees from healthy colonies) or merging (i.e., combining two weak colonies) are expensive interventions as they imply pre-calculated losses of at least one colony, so that they are considered as last resort. Predictive markers can also be used to apply control treatments in time. Treatment can be done in such a way to avoid blind chemical treatment, to prevent resistance development for these acaricides of the mites, and to reduce acaricide residues in honey and bee wax. Late summer is suggested to be the latest for treatment against *V. destructor*, also for merged colonies, to prevent negative effects on the long-lived winter bees (Delaplane and Hood 1997, Currie and Gatien 2006, Van Dooremalen et al. 2012).

Several studies tested the presence and loads of honeybee pathogens or colony health as predictive markers for winter honeybee colony losses (Dainat et al. 2012a, Francis et al. 2013, Braga et al. 2020). *V. destructor* infestation levels in colonies are suggested to be used to anticipate winter losses (Dainat et al. 2012a, Francis et al. 2013, Van Dooremalen et al. 2018). However, the problem of using *V. destructor* infestation levels for early detection of potential losses is that infestation levels are low in the beginning of summer (Sumpter and Martin 2004, Dainat et al. 2012b), and although clinical symptoms are not visible and infestations often remain undetected at low to moderate infestation rates, growth rate of the honeybee colony may be reduced (Rosenkranz et al. 2010). Moreover, estimating *V. destructor* load in colonies in the field is difficult and cumbersome for beekeepers and scientists alike, especially when many colonies should be monitored over time (Lee et al. 2010, Dietemann et al. 2013, Francis et al. 2013). Also, infection with deformed wing virus (DWV) and acute bee paralysis virus (ABPV) in October are suggested as predictors for winter losses (Johnson et al. 2009, Genersch et al. 2010), but that makes them rather late-warning markers. Moreover, costs for beekeeping drastically increase when these types of markers have to be applied at large scale. Therefore, we question whether a simple and fast measure of colony size can be used to predict potential colony losses? This question is central in the development of precision apiculture (Braga et al. 2020). When colony size can be used as early predictor, automatically monitoring hive weight may help beekeepers to prevent losses (Meike and Holst 2015, Meikle et al. 2018).

Some studies present incidental measurements over time of colony size in relation to *V. destructor*. For example, Delaplane and Hood (1997) showed that colonies with high *V. destructor* infestation (in August) had more bees in December than colonies with low infestations, whereas Ostermann and Currie (2004) showed that mite-inoculated colonies with high *V. destructor* infestation had smaller worker populations in August and September than controlled colonies. Only few studies relate colony size to survival during or after winter (Van Dooremalen et al. 2018). Both Genersch et al. (2010) and Dainat et al. (2012b) found that colonies that survived winter were larger in October than colonies that collapsed. These two studies imply that colony size can be a predictive marker of colony losses during winter, but the differences in colony size between infested and uninfested colonies were detected relatively late in the season.

In this paper, we address two questions: (1) whether the decrease in colony size over time during the foraging season is related to *V. destructor* induced losses during winter, and (2) whether the effects of *V. destructor* on colony size can be found early enough in the season, namely before the winter preparation period. To answer the first question, we compared colony size during summer and survival during winter of colonies with high and low *V. destructor* load. We did this by following colonies treated and untreated against *V. destructor* from July until March of the next year. It appeared that most colonies with high *V. destructor* load did not survive until March next year, so we could also use this experiment to answer the second question.

## Materials and Methods

### Experiment

The fieldwork took place in three years between July 2012 and March 2015 at an apiary in Wageningen, the Netherlands. For each year, new colonies (N=20) were used. Colonies were kept in one-story ten-frames wooden hives (inside measures Simplex) and a standard number bees at the start of each year (4 frames of bees in 2012, and 6 frames in 2013 and 2014). Swarming was prevented during the experiment in all colonies. All colonies had continuous access to sugar dough (Apifonda).

For each year, the colonies in the group (N=10) with a low infestation of *V. destructor* (V-colonies) were treated in May/June with oxalic acid spray (30 g oxalic acid dihydrate in 1 L water). At that time, all colonies consisted of bees and open brood only (no capped brood cells). In August, the V-colonies were treated for 6 weeks with Apistan (2 strips per colony). The colonies in the group (N=10) with a high infestation of *V. destructor* (V+ colonies) were neither treated in May/June nor in August. There were no signs that oxalic acids or Apistan resulted in significant bee death neither during nor directly after application (Van Dooremalen et al. 2018).

### Data collection

Colony size was estimated by the percentage cover of the topside of the hive in the first week of each month – when weather permitted (see Van Dooremalen et al. 2018 for the justification of this method). The topside of the hive is what beekeepers see when they open their hive and it is the easiest, less invasive and fastest way to obtain data on colony size, e.g. compared to the more commonly used methods of estimating the number of bees on the different frames or weighing only the bees (Delaplane et al. 2013). A puff of smoke was blown into the hive from below. After a minute, the lid was removed and a photo was taken from the topside. The percentage cover (standardized to one brood box) was calculated based on the number of pixels in the photo, namely the number of pixels of the area covered with bees on top and visible between the frames, divided by the topside of the inner volume of the hive. We used a standard size brood boxes (hives) with 10 frames (Simplex measures). For comparative insight, in this type brood box, when fully occupied with bees (100% top coverage) contains approximately 17.000 (Delaplane et al. 2013). In the first week of next year March, a final check-up of the colonies was performed to determine whether the colony was dead (no living bees and/or queen) or still alive.

The number of phoretic mites was monitored by monthly sampling of 100-200 bees between July and December and counting the number of mites. The worker bees were collected by scraping them off the rim of the brood nest and stored at −20°C until counting. Distributions of age classes are assumed to be similar throughout the brood nest (Van der Steen et al. 2012). Collection of drones or the queen was avoided, but if so not included in the analysis.

All bees in the samples were checked for the number of mites present using a stereo-microscope. Optimally, one would sample at least 300 bees to analyze *V. destructor* population dynamics throughout the whole season with frequent and destructive sampling of bees from healthy colonies (> 10,000 bees) (Dietemann et al. 2013). In our study however, half of the colonies was not healthy and in general our colonies were small (<7000 bees at the start in 2012, estimated based on the assumption that 10% top cover equals 1 frame of bees, and one frame counts for approximately 1700 bees, as estimated by Delaplane et al. 2013). We felt that monthly sampling of 300 bees would reduce the colony size significantly and would therefore interfere with our colony size measurements. We collected per colony per month approximately 12.5 gr bees (∼100 bees) in 2012, 20 gr bees (∼ 160 bees) in 2013, and 15 gr bees (∼120 bees) in 2014. In 2012, we weighed 20 individual bees per colony, resulting in an average body mass of 124.7±1.6 (sd) mg per bee and 8 individuals per gram bees.

To test the reliability of our sampling method we compared the standard error as a percentage of the mean between our dataset based on 100-200 bees per colony and the dataset as presented at Table 1 in Lee et al. (2010) based on 300 bees per colony. The standard error was used (instead of the standard deviation) to correct for the difference in number of colonies sampled between the datasets (10 colonies per group in our dataset, against 30.8±5.7 (sd) colonies per group in Lee et al. 2010). In our dataset, the standard error was 18.6% of the mean in 2012 (100 bees), 15.7% of the mean in 2013 (160 bees) and 18.9% of the mean in 2014 (120 bees), and very similar to the proportion as calculated from Lee et al. (2010), which was 18.5%. We therefore argue that our sampling of 100-200 bees per colony per month was sufficiently reliable to be used to separate low and high *V. destructor* loads.

**Table 1.** Colony survival in March, since start of experiment in July the year before for colonies without acaricide treatment (V+) or with acaricide treatment (V-). Colony # indicates that number of starting colonies.

|  | V+ |  | V- |  |
| --- | --- | --- | --- | --- |
| Year | Colony # | Survival | Colony # | Survival |
| 2012 | 10 | 0 | 8 | 8 |
| 2013 | 10 | 2 | 10 | 10 |
| 2014 | 10 | 3 | 10 | 10 |

### Statistics

General Mixed Linear Models were used to test the dependent variables the number of mites per gram bees (Ln-transformed +0.01) and the colony size as a function of the acaricide treatment (fixed factor) or survival in March (fixed factor), year, and sampling month (repeated, colony as subject). The model with the lowest AIC in relation to the covariance structure was used.

The use of the percentage cover of the topside of the hive to estimate colony size is sensitive to differences in ambient temperature. To draw any conclusion about using colony size as a reliable predictive marker for winter loss, one should compare the sizes of the colonies under similar temperature conditions. Therefore we tested the relation between colony size (Ln-transformed) and average daily temperature (Ln-transformed). We used a Linear Mixed Model with year as fixed factor and average daily temperature as covariate.

In all models, Sidak posthoc tests for pair wise comparison were used to test for differences between groups within each month. Assumptions for normality were met in all tests. Unless explained differently, means are presented together with their standard error.

## Results

Most of the V+ colonies that were not treated against *V. destructor* died during the winter, whereas all V-colonies survived until March of the next year (Table 1). The number of mites per gram bees was lower in 2012 compared to the years 2013 and 2014 (not different), but in all years the number of mites per gram bees in V+ colonies increased towards the end of the year (Table 2, Fig. 1). From September onwards, V+ colonies showed higher infestation levels compared to V-colonies. We noticed that many of the colonies not treated with acaricides did not show any infestation in July or August (Table 3).

**Table 2.**
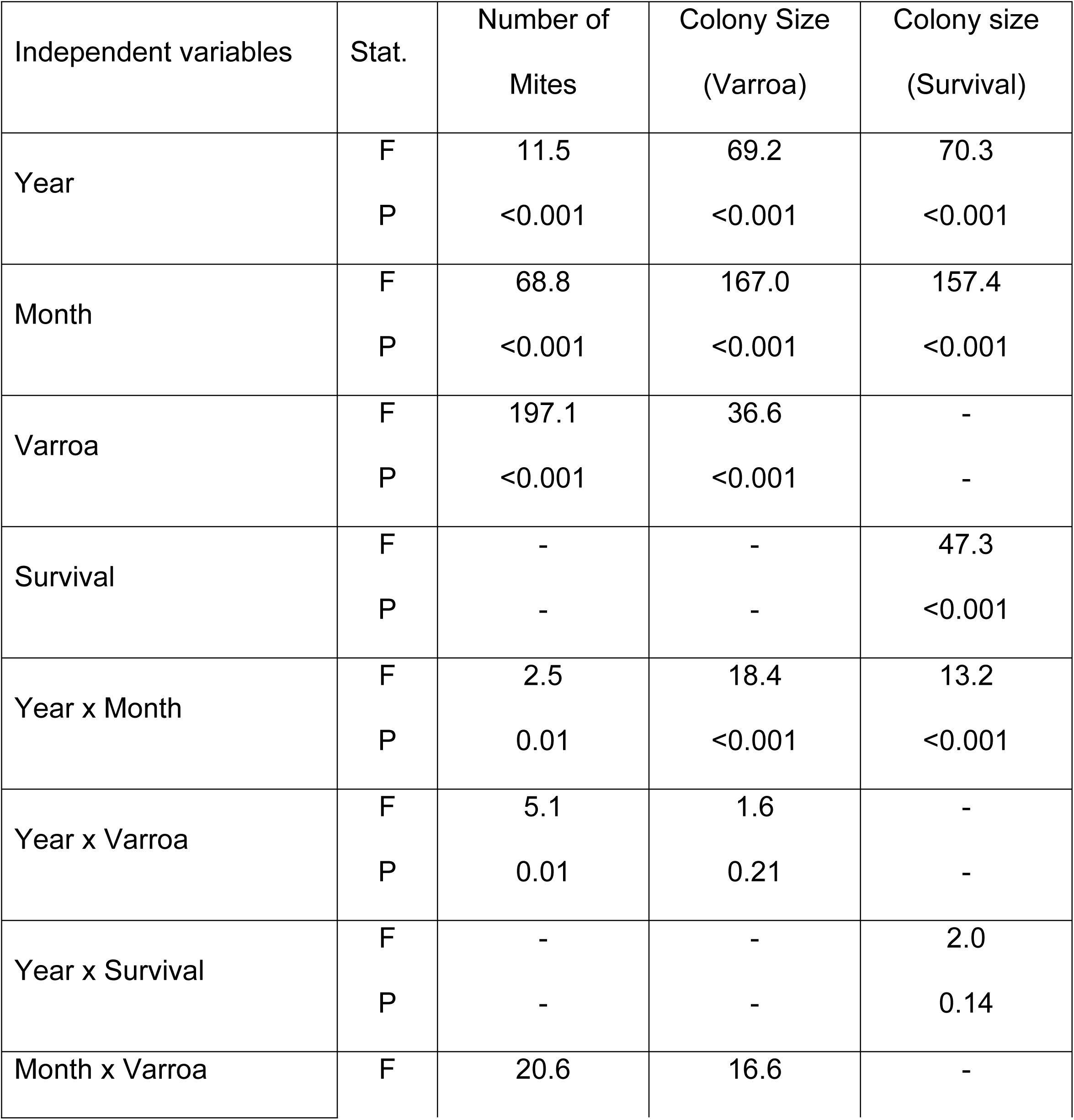
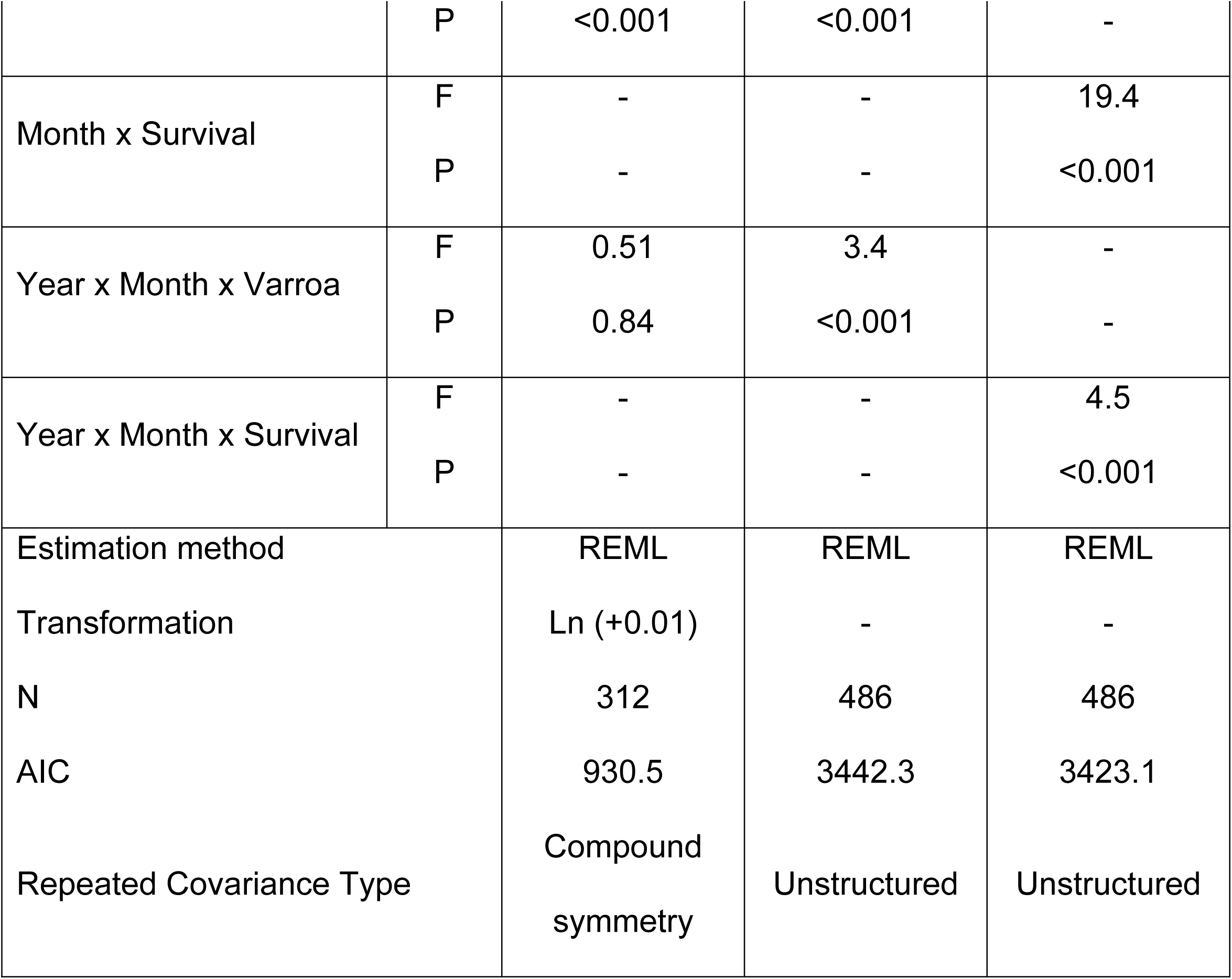
Results of the linear mixed models for the effects *Varroa destructor* (Varroa), Year and Month, including their interaction on the number of mites per gram bees and the colony size (% top cover). For each factor in the model, the F- and P-value are given. For each model, we give the applied method of estimation (REML= Restricted Maximum Likelihood), whether we Ln-transformed the dependent variable (Ln), the sample size (N), the value of the Aikaike’s Information Criterion (AIC) and the Repeated Covariance Type.

**Table 3.** Percentage of samples with no mites in the group of colonies not treated with acaricides (V+) for the different years and months.

|  | Year |  |  |
| --- | --- | --- | --- |
| Month | 2012 | 2013 | 2014 |
| Jul | 40 | 90 | 80 |
| Aug | 20 | 80 | 40 |
| Sep | 0 | 10 | 10 |
| Oct | 0 | 0 | 10 |
| Nov | 0 | 0 | 0 |
| Dec | 0 | 0 | 0 |

**Fig. 1.**
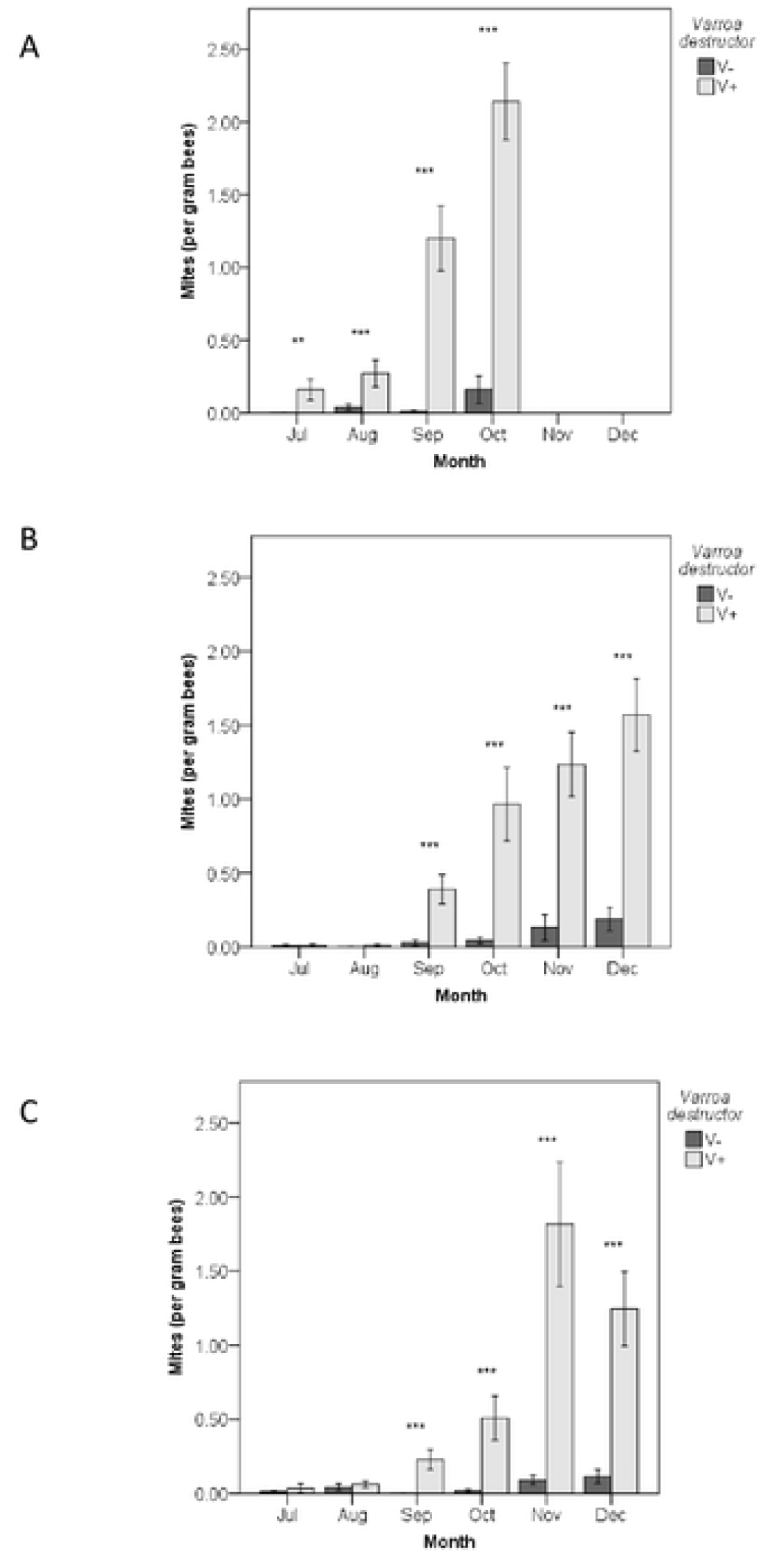
Number of *V. destructor* mites per gram bees per month in 2012 (a), 2013 (b), and 2014 (c) for colonies without acaricide treatment (V+, dark grey bars) or with acaricide treatment (V-, light grey bars). Asterisks show differences between V- and V+ (*P<0.05;** P<0.01; *P<0.001), where the data for the statistical test were Ln-transformed (+0.0005).

Colony size differed per year, month, and *V. destructor* infestation (Table 2, Fig. 2). However, differences in colony size due to *V. destructor* infestation did not in general appear before December (Sidak post hoc test Varroa x Month). In one year (2014) differences due to *V. destructor* levels appeared a little earlier in the year, in November (Fig. 2c). Differences in colony size based on survival (dead/alive in March) did also not appear during summer (Table 2, Fig. 2). In October, the earliest difference in size was observed between the colonies that did not survive until March and the ones that did (2014, Fig. 2f).

**Fig. 2.**
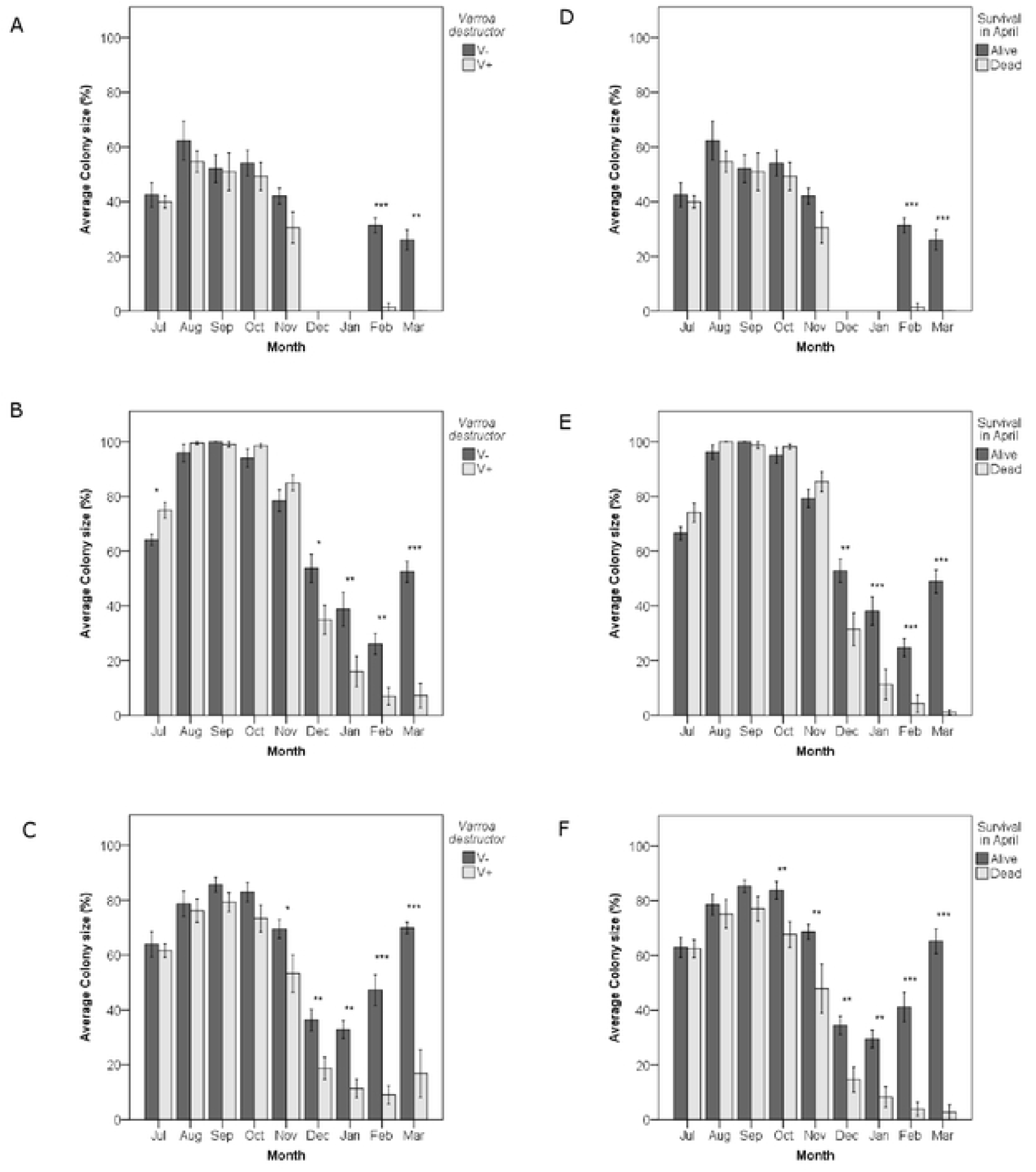
Average colony size (%, Van Dooremalen et al. 2018) per month for 2012 (a,d), 2013 (b,e), and 2014 (c,f) for colonies with (V-, dark grey bars) or without (V+, light grey bars) acaricide treatment (a-c) or colonies alive (dark grey bars) or dead (light grey bars) in March (e-f) Asterisks show differences between series (*P<0.05; **P<0.01; ***P<0.001).

In 2012, we started with 25% smaller colonies compared to 2013 and 2014 (not different). Colony sizes were largest during summer and smallest in winter. This general pattern of colony size over the experimental months was positively related to the average daily ambient temperature in the same months (Year F_2,19_=0.3, P=0.72; Average daily temperature per month F_1,19_=41.0, P<0.001; interaction F_2,19_=0.7, P=0.50; both Ln-transformed; Fig 3). The lack of an interactive effect between month and temperature on the colony size in our analysis justifies comparing colony size under similar temperature conditions, which allows us to draw the conclusion about the use of colony size.

**Fig. 3.**
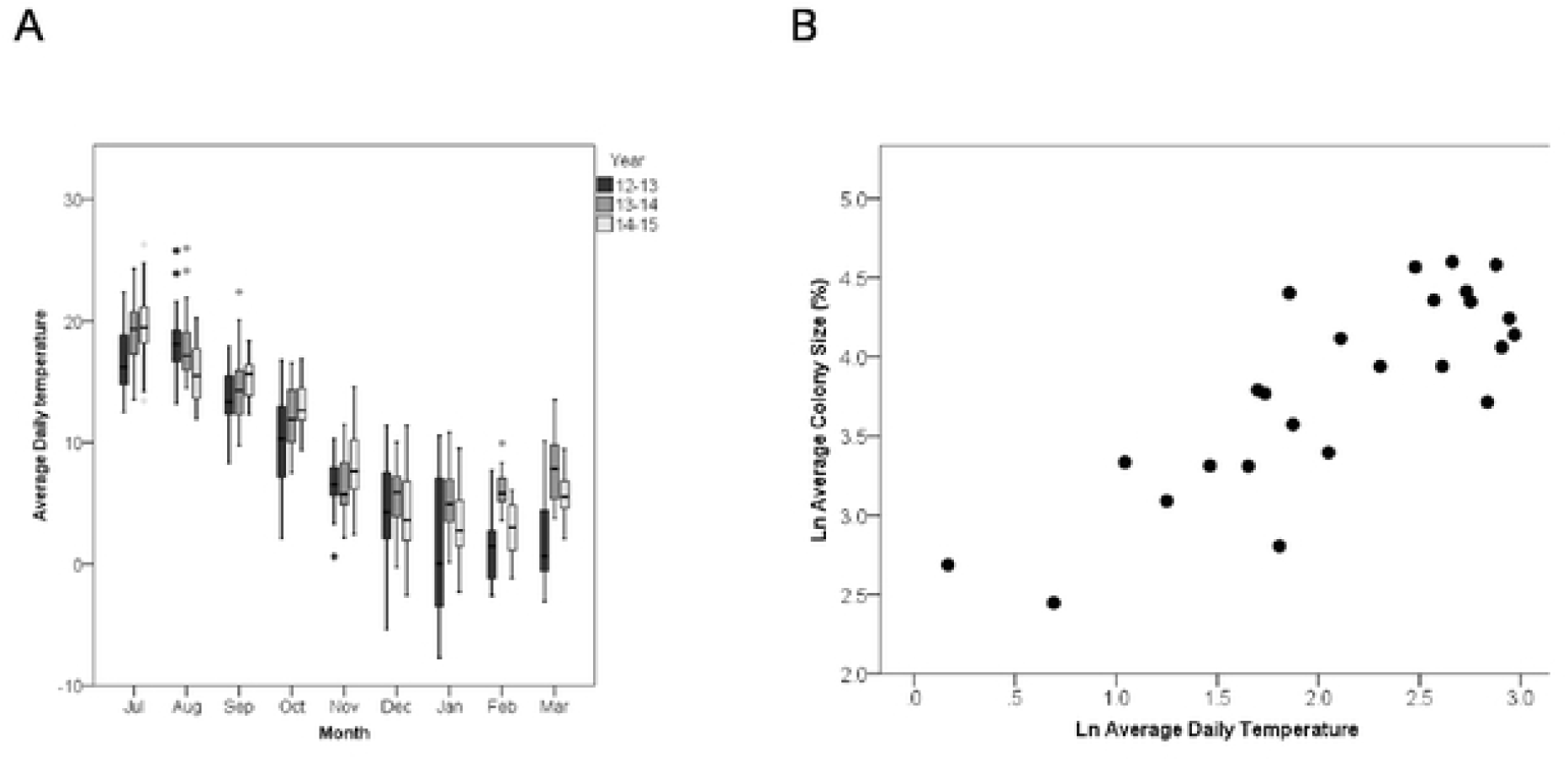
Average daily temperature (°C) per month for 2012, 2013, and 2014 (a) 2 and its relationship with the average colony size per month (%) (b). In panel b, the Ln-transformed data are shown.

## Discussion

In this paper, we investigate whether colony size can be used as early warning predictor for colony losses during winter that are caused by the parasitic mite *V. destructor*. In our experiment covering three years, we found that the colonies with high load of *V. destructor* had a much higher probability of mortality before next spring than the colonies with low load. The differences in colony size between the colonies with low and high load of *V. destructor* were only visible after November, even in years when differences in mite levels already occurred in July. Mere visual inspections of the colony sizes will therefore be insufficient as predictive marker to prevent winter losses; late autumn is considered to be too late for beekeepers to strengthen colonies in preparation of winter. We acknowledge that due to different climatic conditions, the honeybee season can end earlier or later in different regions, but these differences do not alter our conclusion that differences in colony size are detectable only late in the season and hence cannot be used as early warning for winter losses. Previous studies looking at differences in colony size as result of *V. destructor* support our findings that differences in colony size are visible too late, although these studies did not investigate a time series of colony sizes as we did: smaller sizes in colonies with high *V. destructor* load in August and September (Ostermann and Currie 2004), in October (Genersch et al. 2010, Dainat et al. 2012b) and in December (Delaplane and Hood 1997) than control colonies.

The lack of differences in colony size during summer can be explained by a larger effect of *V. destructor* on the condition and life span of individual bees than on the number of bees in the colony *per se* (Rosenkranz et al. 2010, Van Dooremalen et al. 2012, 2013, 2018, Blanken et al. 2015). No effects were namely found of different levels of *V. destructor* infestation on the number of brood cells in colonies (Van Dooremalen et al. 2012), i.e. the future work force. Regardless of the absence of an early effect of *V. destructor* on colony size, we cannot reject the hypothesis that colony size may still be a valid marker to early predict colony losses in response to other stressors – especially those which (in)directly impact the number of bees in the colony (Perry et al. 2015), such as pesticides (modelling study in Henry et al. 2012).

In our study, infestation levels showed already differences between groups from July (2012) or September (2013 and 2014), but the numbers of (phoretic) *V. destructor* mites in these months were very low and many colonies in the untreated group (V+) showed no mites yet (40-90% of the colonies in July; 20-80% of the colonies in August). We assume that most of the mites will be reproducing in the brood cells during this period and only very small numbers will be in the phoretic phase (Rosenkranz et al. 2010, Van Dooremalen et al. 2012, Dietemann et al. 2013). Due to the large variation in infestation levels and mite population growth patterns between years and the low phoretic mite levels during summer, our study indicates that the infestation level is also not very useful as a reliable predictive marker in an easy, quick and strait forward way during summer. Similar drawbacks are expected for other mite-related parameters such as visual inspection and counting the number of DWV symptoms.

The devastating effect of *V. destructor* infestation on colony survival is well known. Fries et al. (2003) and Rosenkranz et al. (2006, as cited in Rosenkranz et al. 2010) found that untreated colonies which exceeded an infestation rate of 30% in the adult bees during summer did not survive the following winter. Additionally, levels above 6% showed more than 10% losses (Genersch et al. 2010, Rozenkranz et al. 2010). In our study, we found lower thresholds: a mite infestation on adult bees (V+ group) in September as low as 3% resulted in 70% loss (2014), 5% resulted in 80% loss (2013), while 15% resulted in 100% loss (2012) of colonies during the winter. Interestingly, even with such high losses during winter among the colonies with the highest mite infestation, no difference in colony size was observed during summer and autumn. This means that beekeepers that are able to partly reduce mite infestation will be even much less likely to find visual differences in colony size, but probably still have high chances of winter losses.

There is clearly a need to search for colony traits as predictive markers, rather than traits of individual bees (e.g. pathogen loads; Dainat et al. 2012a, Francis et al. 2013), because measuring individual traits such as pathogen load in the field is time- and money-consuming, especially when beekeepers have many colonies that should be monitored over time. The way to develop such methods is to test the relative and interactive effects of multiple stressors in experiments on winter survival of colonies, following colony traits that indicate the functioning and condition of colonies from spring until the next spring (Van Dooremalen et al. 2018). Braga et al. (2020) developed a classification algorithm based on a supervised machine learning approach to estimate the health status of colonies and to indicate an imminent collapsing state to beekeepers. Their promising results suggest a high precision classification model, which can be useful to self-predict healthy, unhealthy, and collapsing bee colony health states. This method and others in precision apiculture (Henry et al. 2019, Catania and Vallone 2020) provide promising, non-invasive ways to measure colony health. Experiments to test the potential of colony traits for early prediction of winter losses to feed these approached are still needed, including the development of devices or tools that allow easy and quick measurements of these colony traits.

## Acknowledgements

The project was funded by the Ministry of Agriculture, Nature Conservation and Food Quality (LNV) of the Netherlands (BO-06-012) and by the European Union (National Honey Program NL08/2.1), year 2012/2013.

